# Tree-structured topic modelling of single-cell gene expression data uncovers hierarchical relationships between immune cell types

**DOI:** 10.1101/2023.11.06.565879

**Authors:** Patricia E. Ye, Yichen Zhang, Ramon I. Klein Geltink, Yongjin P. Park

## Abstract

Immune cells undergo a series of differentiation steps following a lineage-tree structure stemming from hematopoietic stem cells. During differentiation of immune cells in both homeostasis and pathological processes, many gene regulatory mechanisms are shared by fully differentiated immune cell sub-types. In order to characterize these features quantitatively, we propose `LaRCH`, a tree-structured embedded topic model. In this model, single-cell gene expression profiles are represented by a mixture of topics consisting of latent features that follow an underlying tree structure, mirroring that of cellular differentiation–nested cluster structures. We present findings of our model trained on simulated single-cell RNA sequencing (scRNA-seq) based on cell-sorted bulk RNA-seq data as well as on a scRNA-seq dataset of over 1.2 million cells from healthy individuals and individuals diagnosed with systemic lupus erythematosus (SLE). The cellular topic profiles estimated by our model markedly improve clustering accuracy over traditional latent variable models and illustrate transcriptomic differences between SLE phenotypes, revealing a pivotal role of multiple immune cell types in disease progression and relapse. Ultimately, `LaRCH` captures the hierarchical context between cellular subtypes by simultaneously identifying shared and distinct latent features amongst subsets of heterogeneous samples of cells.

## 1 Introduction

### Background

Immune responses rely on the coordinated efforts of many different cell types with specific roles in order to respond to pathogens and protect from disease. Most immune cell subsets are derived from pluripotent hematopoietic stem cells and follow a process consisting of many consecutive differentiation steps in order to develop into their mature state [1]. Immune cell differentiation is driven by a number of factors, including the presence of microbes, metabolites, and subset-specific regulatory cytokines in the extracellular environment as well as the expression of transcription factors (TFs) regulated in part by epigenetic mechanisms. A lineage tree is often used to represent this differentiation process as certain cell types share lineage paths with others [2, 3]. Aberrant regulation of immune cell function or altered subset composition results in the development of many immunological diseases, such as systemic lupus erythematosus (SLE) [4]. SLE is an autoimmune disease classified by persistent inflammation in vital organs, fever, and fatigue. The pathogenesis of SLE is an area of ongoing investigation, but one known driver of SLE is the presence of autoreactive T and B cells, resulting in persistent inflammatory responses in the absence of pathogens [4]. A better understanding of the cellular makeup of immune cells in the body provides further insight into causal mechanisms in autoimmune disorders.

With the recent advancements of high-throughput single-cell sequencing technologies, including single-cell RNA sequencing (scRNA-seq) and single-cell Assay for Transposase-Accessible Chromatin with sequencing (scATAC-seq), knowledge of the molecular profiles of specific cell types has expanded. It has been suggested to use a tree-based reference structure to represent cell types as opposed to an atlas or periodic table-like classification system in order to better capture relationships between cell types [5]. However, unsupervised machine-learning approaches are more frequently used to identify cell clusters that constitute an essential basis for unbiased cell-type annotation results.

### Autoencoder models

More recently, deep-learning-based approaches have gained popularity, including scAlign [6], scETM [7], and scVI [8]. These methods borrow from neural network architectures originally invented for natural language processing [9] and show state-of-the-art performance, generally outperforming traditional factorization-based methods. The basic idea of the autoencoder models is to encode high-dimensional single-cell gene expression vectors onto a smaller latent space. A common drawback of these methods, keeping in mind the benefit of a reference cell tree, is that features within the latent space share a flat representation, not assuming nested structures between cell states/topics. Moreover, neural network model parameters often create another challenge in interpretation, often demanding *post-hoc* analysis.

### Post-hoc methods

Methods to build tree-structured relationships of single-cell sequencing data are emerging. treeArches [10] uses single-cell data to construct and extend a cell reference atlas and corresponding hierarchical classifier, while cellTree [11] produces tree-structured relationship hierarchies between cells. These methods construct their relevant cellular tree after estimating their latent representations rather than learning the cellular hierarchies simultaneously with their latent representations.

### Probabilistic methods

Hierarchical extensions to topic modelling have been proposed, such as hLDA [12], which uses a Latent Dirichlet Allocation (LDA) to represent topics in a hierarchical structure informed by a nested Chinese restaurant process prior, and TSNTM [13], a tree-structured neural topic model that builds on a variational autoencoder framework. These methods rely on Markov Chain Monte Carlo algorithms, simulating models over super-exponential space. Despite theoretical guarantees, we found these methods were not suitable for dealing with scRNA-seq data of millions of cells over tens of thousands of features.

### Our contribution

We propose a tree-structured <u>La</u>tent <u>R</u>epresentation of <u>C</u>ellular <u>H</u>ierarchies (`LaRCH` for short), an extension of the embedded topic model framework [14] with a built-in underlying tree-structure informing a basis for hierarchical cellular relationships. By enforcing a structured topic model, we are able to cluster and classify cells from scRNA-seq data while uncovering shared latent features between cell types in a finite search space. Using variational inference for training results in a computationally feasible model for large high-dimensional gene expression data. The application of `LaRCH` on simulated and real data sets shows the effectiveness of a structured topic model for cell-type discrimination. Training our model on scRNA-seq data of peripheral blood mononuclear cells (PBMCs) from individuals with varying SLE disease conditions shows pathological differences in immune cell composition and specific genomic latent features.

## 2 Results

### 2.1 LaRCH model overview

`LaRCH` builds upon an embedded topic model (ETM) framework [14] with an additional tree-node layer that aggregates sparse gene expression effects informed by a Bayesian spike-and-slab prior [15]. Within a typical ETM, cells are represented as a mixture of latent topics each describing a gene expression profile of raw scRNA-seq count data. `LaRCH` introduces an additional layer of latent nodes that lie within a perfect binary tree structure (Fig.1a). As a result, cells are viewed as a mixture of latent topics, each corresponding to a summation of tree nodes along a path from the root node to a given leaf node of the underlying tree. Rather than choosing the number of latent topics to represent the data, we choose the depth of the latent node tree *D*, so the resulting number of tree nodes is 2^*D*^*−* 1, and the number of latent topics to represent the cells is 2^*D−*1^. The `LaRCH` model is comprised of an encoder and decoder component. The encoder transforms raw count data into the latent topic space and inferred topic proportions through a neural net. The decoder contains a gene embedding matrix *β* for each tree node as model parameters, following a spike-and-slab prior distribution to induce sparsity. The learned embedding matrix and inferred topic proportions are passed to a generalized linear model (GLM) to estimate the cell-type-specific gene frequencies *ρ*. Likelihood values for expected gene counts are computed from a multinomial distribution with *ρ* as the distribution parameter.

**Figure 1:**
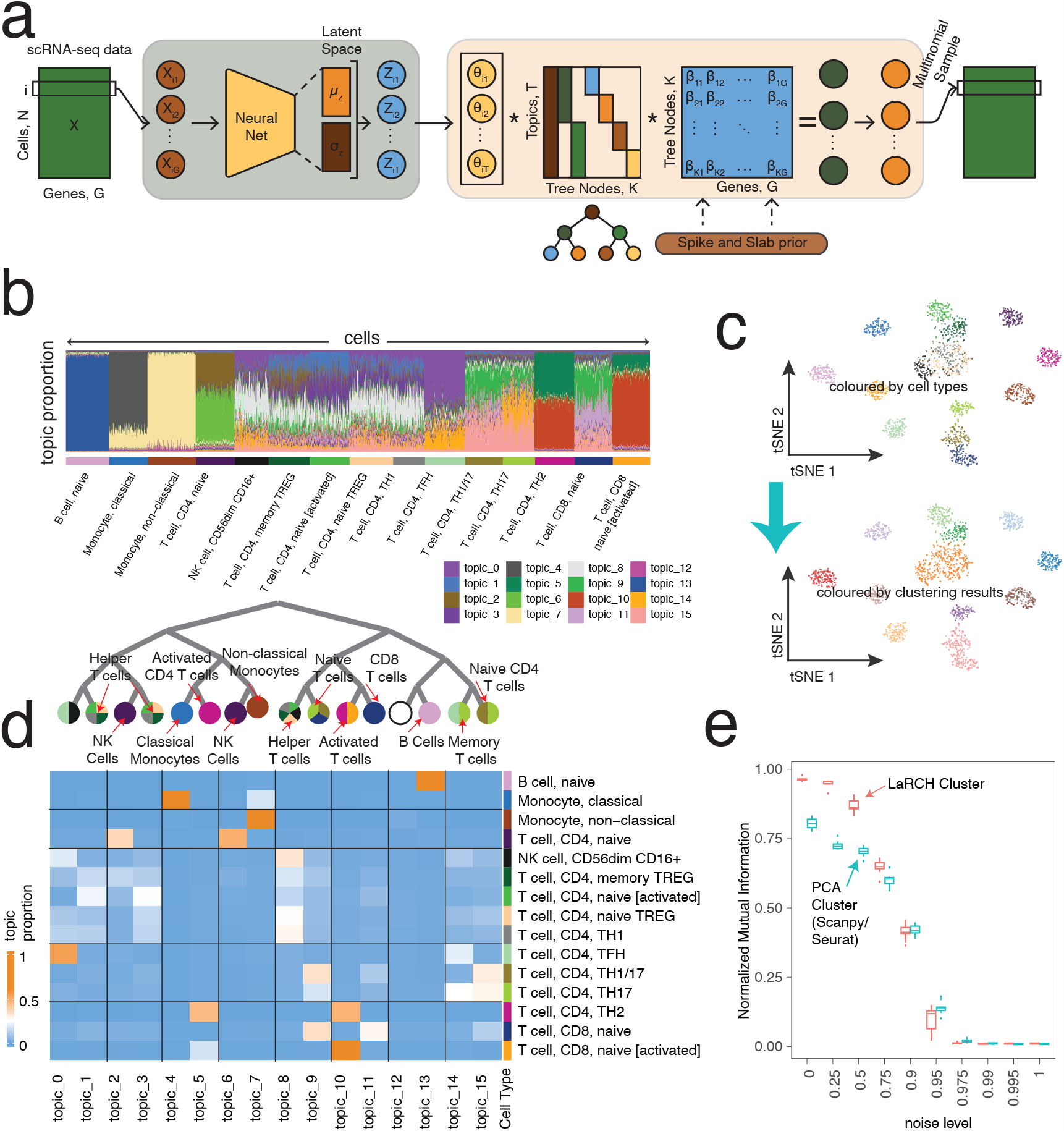
LaRCH effectively identifies cell types and uncovers their relationships in realistic simulation data. a) Overview of the `LaRCH` model showing both the encoder and decoder components. b) Estimated topic proportions across 2000 simulated immune cells. c) *t*-distributed Stochastic Neighbourhood Embedding (`tSNE`) projection using latent topic features as input. Coloured by cell type (top panel) and Louvain cluster (bottom panel). d) Visualization of tree-structured relationship between cell types and mean latent topic proportion profile across each cell subtype. e) Bar plot showing normalized mutual information between clusters generated from latent features or principal components and true cell type for simulated data sets across varying noise levels.

### 2.2 LaRCH learns cell types and hierarchical relationships in simulated data sets

We simulated single-cell gene expression count data of immune cells informed by cell-type sorted bulk RNA-seq gene expression profiles from the Database of Immune Cell Expression (DICE) [16]. From a `LaRCH` topic model trained on a simulated dataset of 2000 cells, noise proportion 0.5, and 2500 reads per cell, we see that our model is able to recover the simulated cell types and represent each cell type with a unique topic profile (Fig.1b,d). Similar cell types share topics, seen most prominently amongst helper CD4+ T cell subsets which contain mixtures of topics 1, 3, and 8 in different proportions (Fig.1d).

Using the topic representation, we reconstruct the underlying tree structure of cell types (Fig.1d). Cell types that are more similar in function and share canonical lineage paths tend to lie within closer points in the tree, such as topics 8 through 11, which solely represent T cell subtypes. More distinct cell types are contained within unique branches of the tree, such as B cells which are the only cell type represented in the branch containing topics 12 and 13, or the monocyte subtypes exclusively represented by topics 4 and 7, which share three nodes within their paths.

Performing tSNE dimensionality reduction on the latent representations of cells shows the distance between cell types in the latent space (Fig.1c). Using the Louvain method of community detection [17] on K-nearest neighbour graphs (*k* = 10) using the latent topic features to calculate distance results in clusters that demonstrate high congruence between cell types and clusters, missing only some granularity (Fig.1c). We simulated datasets at varying noise levels (10 each) and compared the accuracy of clusters in both the latent topic space and principal component space (Fig.1e). Clusters generated from the latent topic space show higher normalized mutual information with the true cell-types up to reasonable noise levels in the simulated data.

### 2.3 LaRCH effectively models real scRNA-seq data from an SLE patient study

We trained our topic model on the scRNA-seq dataset from Perez *et al*.[18]. This dataset consists of single-cell transcriptome data of 1.2 million peripheral blood mononuclear cells (PBMCs) from 99 healthy control individuals and 162 individuals with SLE. Estimated topic proportion profiles show distinct groups of cells within the latent space that align closely with the cell types determined in the original Perez *et al*. paper [18] (Fig.2a,e). The topic representation of cells is able to pick up more granularity that potentially describes further distinct cell types, namely in the cells labelled as ‘classical monocytes’ where there are two distinct groups in the topic space primarily represented by either topic 10 or topic 11, suggesting classical monocyte-labeled subsets could be composed of subsets with differing cell function.

**Figure 2:**
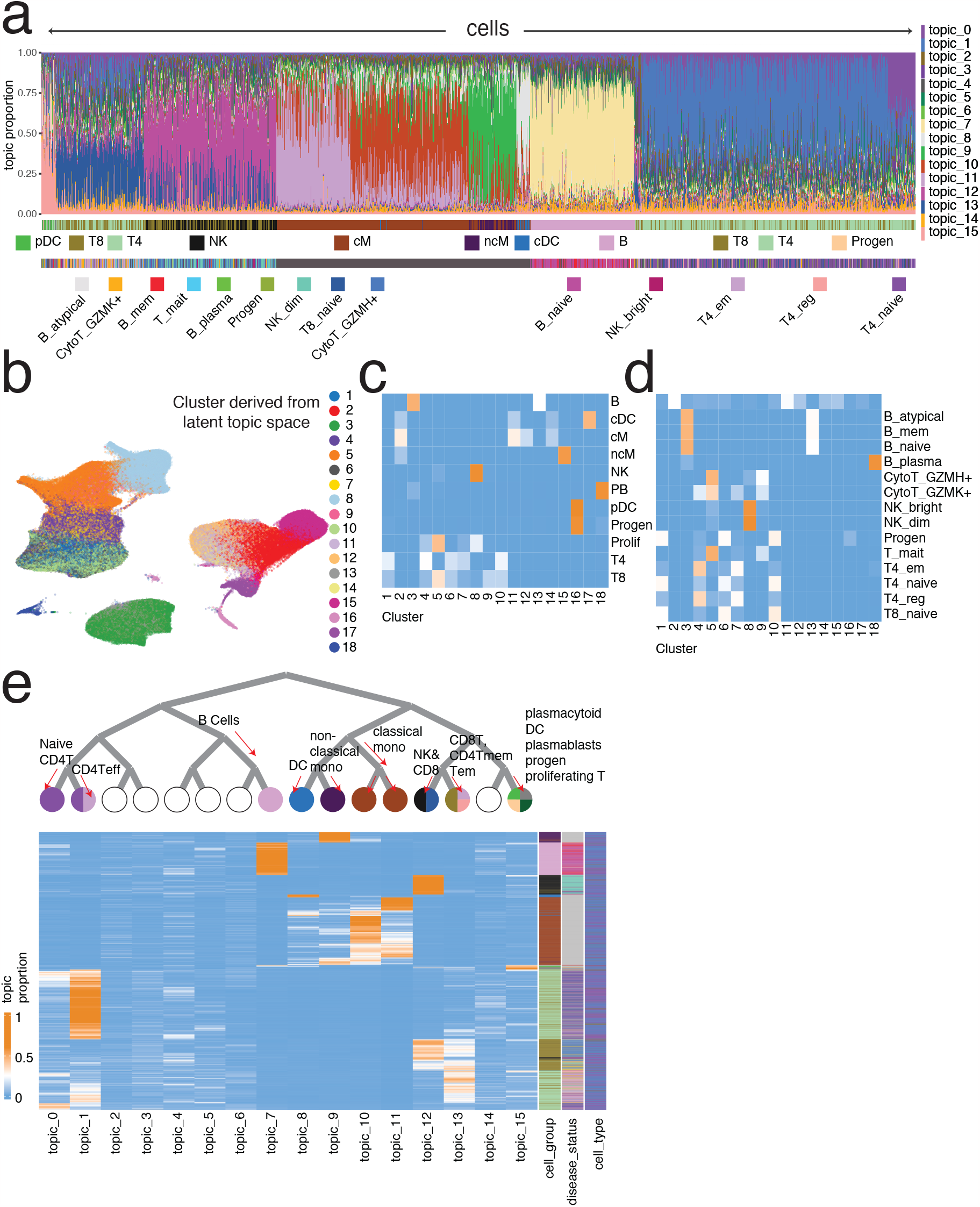
LaRCH discriminates cell types and constructs a biologically relevant cellular hierarchy in a real scRNA-seq dataset. a) Estimated topic proportions of a subset of 10,000 PBMCs from healthy individuals and individuals with SLE annotated by proposed cell group and type. b) UMAP projection from Perez *et al*. [18] coloured by Louvain cluster derived from latent topic space. c) Similarity matrix between Louvain cluster and annotated cell groups. d) Similarity matrix between Louvain cluster and annotated cell types. e) Visualization of reconstructed cellular hierarchy relationships with heatmap showing estimated topic proportions of a subset of 10,000 cells.

Similar to that from simulated data, the constructed tree structure from the real data set aligns with the canonical understanding of immune cell differentiation (Fig.2e). B cells lie within their own distinct branch of the tree. Myeloid cells, including monocytes and dendritic cells are contained within the tree branch with topics 8 through 11. Topic 15 captures a number of intermediate immune cell-states, including progenitor and proliferating lymphoid cells, proliferating dendritic cells, and plasmablasts. Naive CD4+ T cells are primarily represented by topic 0 and 1, while the remaining CD4+ subtypes are represented by topic 13 along with non-naive CD8+ T cells. NK cells and naive CD8+ T cells are found in topic 12.

Louvain clustering [17] using *k* = 30 nearest neighbours in the latent topic space results in clustering that aligns with cell groups described by Perez *et al*. [18] (Fig.2b,c), *but quickly loses the ability to clearly distinguish cell the more specific cell types (Fig.2d). Since the original cell types are assigned from Louvain clustering on UMAP reduced dimensions and canonical marker gene expression, it is unclear as to which clustering method produces higher-quality immune function-relevant cluster assignments*.

### 2.4 Stratified analysis shows SLE disease status dependent latent topic profiles

*From the provided metadata of the scRNA-seq dataset in Perez et al*. [18], we analyzed our trained model in a disease-stratified fashion. Using the same Louvain clusters determined in the previous section (Fig.2b), we found differentially represented clusters depending on SLE disease status (Fig.3b). Clusters 1, 6, 7, 9 and 11 capture primarily cells from healthy individuals or individuals with managed SLE. Cells in clusters 2 and 12 make up a significantly greater portion of cells from individuals with any level of SLE compared to healthy individuals. Some clusters make up a greater portion of cells from individuals with active disease phenotypes of SLE (treated vs. flare), such as clusters 4 and 5. Finally, cluster 10 appears in each condition except for individuals with managed SLE. The differences in cluster representation suggest that the cellular makeup of PBMCs is associated with disease status. To understand how each cluster relates to the estimated latent topic representation of cells, we constructed the average topic profile for cells in each cluster (Fig.3a). From this, we see that many clusters that are differentially represented across disease conditions actually share many features, suggesting that cells in different clusters may correspond to the same general cell type, but have some minute differences in gene expression. For example, clusters 4 and 7 both have strong representations by topics 1 and 13 but cells in cluster 7 have a higher representation in topic 0. Cells in these clusters are shown to be most closely aligned with effector memory and regulatory CD4+ T cells (Fig.2c,d), yet are distinctly different in their latent topic profiles.

**Figure 3:**
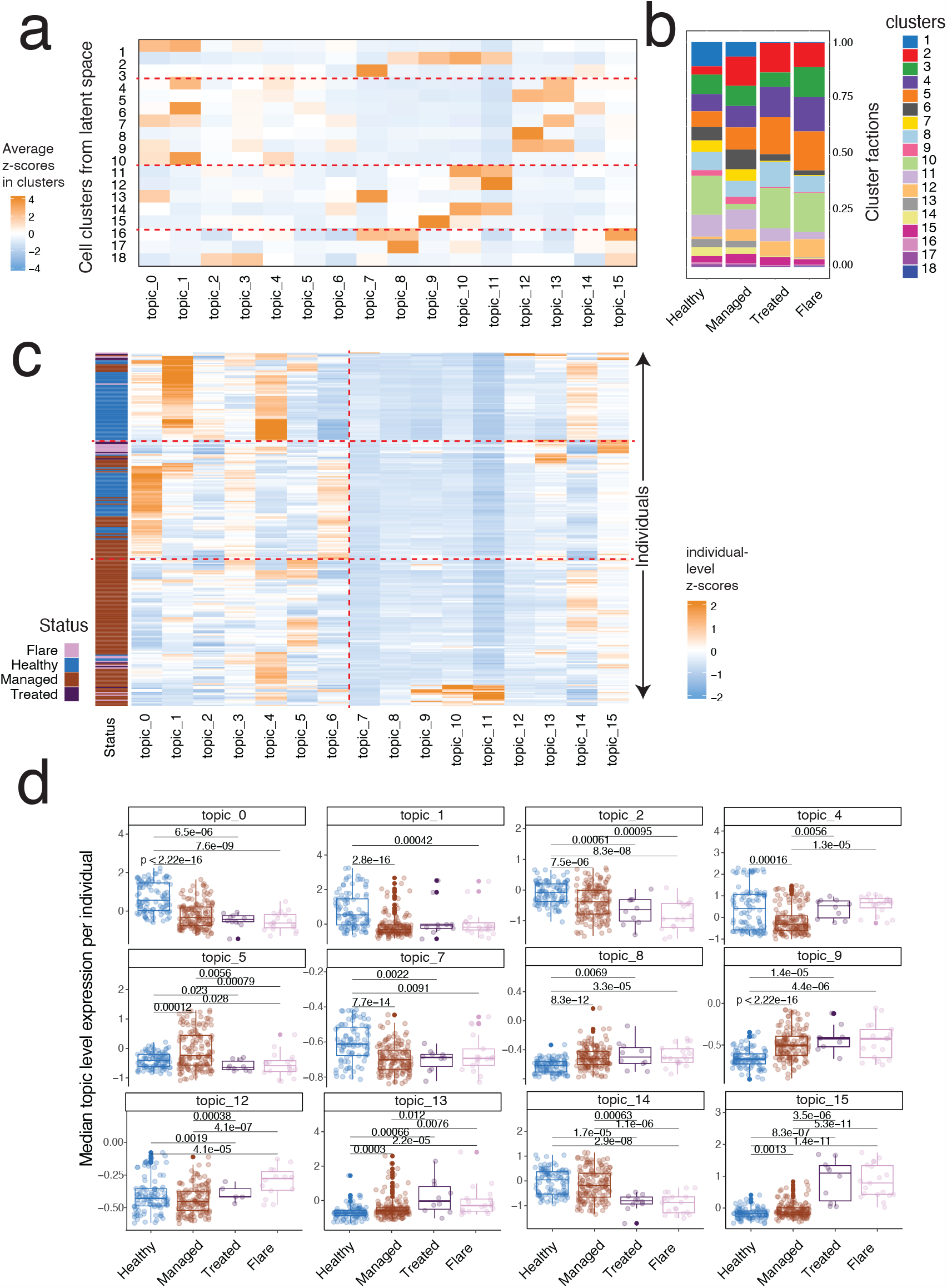
Cell type composition and latent topic values are SLE disease status dependent. a) Median latent values across cells in each Louvain cluster. b) Cluster proportion breakdown by SLE disease status. c) Median latent values per individual in the SLE study, annotated by disease status. d) Boxplot showing individual level median latent values of a selection of topics. Annotated p-values from Wilcoxon rank-sum tests.

To better understand differences in topic representation as an effect of disease phenotype, we constructed the median topic profile for each individual included in the study (Fig.3c). From the latent profiles, we see a large variance in topic profiles across individuals, yet individuals with similar profiles tend to share the same disease status. Wilcoxon rank-sum tests were performed on median latent topic values of individuals between disease status groups (Fig.3d). Of note, topic 0, which separates cells found in cluster 7 from those in cluster 4, is significantly differentially represented in healthy individuals compared to individuals with SLE of any status. Also healthy individuals and those with managed SLE have a significantly higher level of topic 14 than those with an active SLE status, while the inverse is true for topic 15, suggesting a higher prevalence of intermediate immune cell states in individuals with active SLE while healthy and managed individuals have a higher prevalence of B cells and certain naive T lymphocytes (Fig.2c,d, Fig.3a). Showing the differences in topic levels across disease statuses informs our investigation into the differentially regulated biological gene sets suggested by the latent-node level gene embeddings of our model.

### 2.5 Functional cell features described by node-level model parameters

We further investigated how many genes effectively define cell-type-specific disease mechanisms using learned node-level gene vectors from `LaRCH`, namely 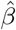 parameters before the tree-based topic-level aggregation. We assumed that Bayesian variable selection prior can put zero values to unnecessary (or statistically weak) associations for irrelevant genes. Counting only the number of genes significantly deviating from zero, we counted the number of genes needed for each tree node and level (Fig. 4). For brevity, we clustered genes into 25 distinctive gene modules by applying the Louvain clustering method on the 10-nearest neighbour gene-gene interaction graph (Fig.4a-b). Bayesian spike-and-slap prior ensured that the node-level gene programs are determined by a subset of genes between 50 to 1800 genes, with a median number of 429. Albeit, we found more genes on nodes 5 (cluster 24) and 7 (cluster 19), which correspond to T- and B-cell groups, respectively.

**Figure 4:**
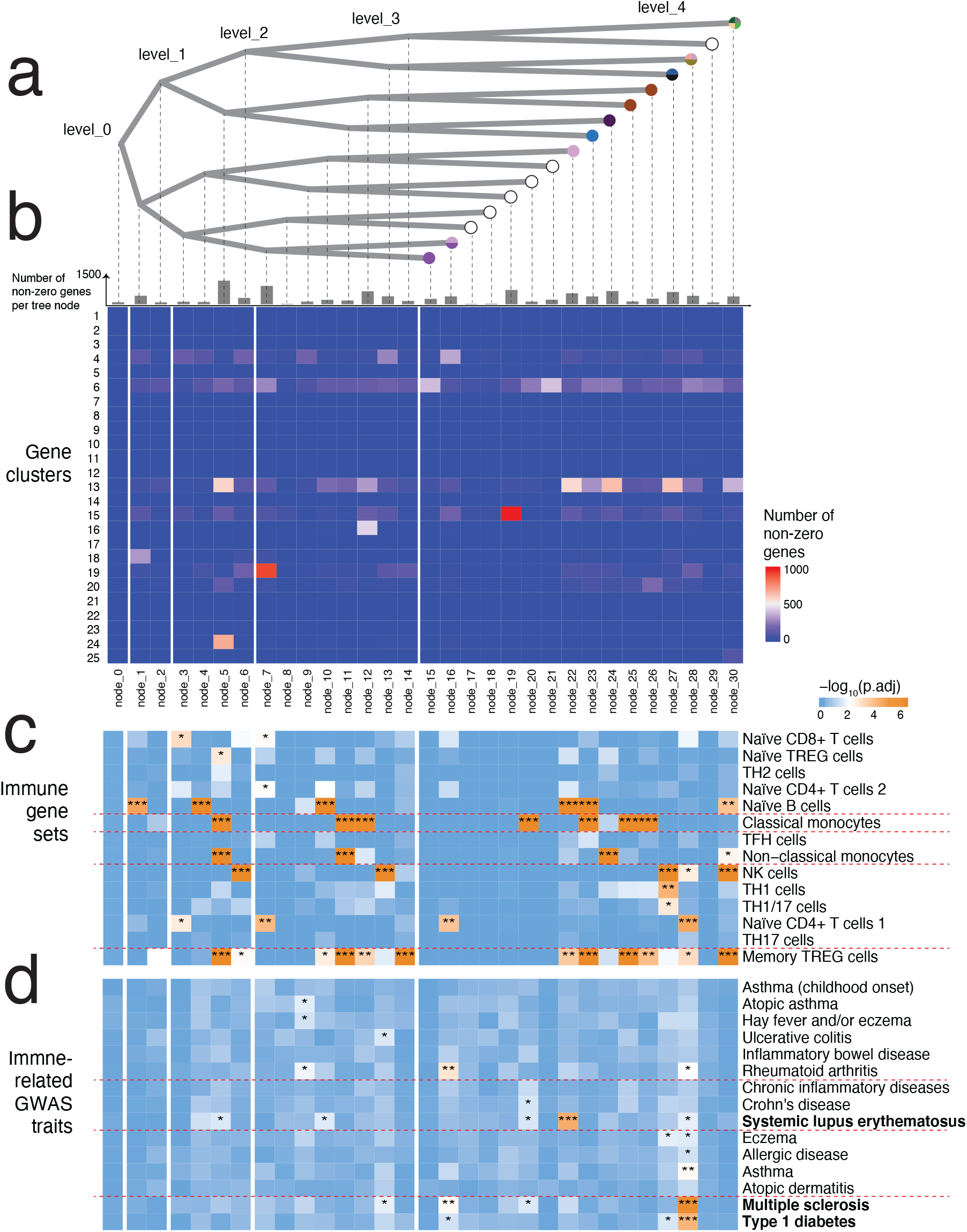
Node-level model parameters provide valuable insight into differences in cellular function implied by latent topic values. a) Tree-structure representation corresponding to node-level features. Leaf nodes are annotated with immune cell types described in Fig.2e. b) Heatmap showing the overlap in gene sets between Louvain gene clusters and significantly enriched genes per node. c) Node level enrichment of immune cell-type specific gene sets. d) Node level enrichment of GWAS gene sets relating to immunological disease.

Enrichment analysis of cell-type specific differentially expressed gene sets based on gene embedding values allows for further annotation of cell-type relationships to latent topics (Fig.4c). We see that B cell-related genes are enriched at the early stages of the node tree, suggesting a distinct model lineage that these cells follow. We also see an enrichment of B cell-related genes in node 23, along with classical monocyte and memory regulatory T cell-related genes. In Fig.2e, topic 8 (made up of nodes 0, 2, 5, 11, and 23) was determined to correspond to dendritic cells, which canonically share lineage paths with both monocytes and lymphoid cells [1]. Topic 1 (made up of nodes 0, 1, 3, 7, and 16), which is significantly greater in healthy individuals (Fig.3d), is enriched in genes relating to naive T cells, suggesting a loss of naive cells and concomitant increase in activated T cells in individuals with SLE. Gene sets of less prevalent sub-types, such as Th17 and Th2 helper T cell subsets, are not found to be significantly enriched in any node as they are difficult to distinguish amidst a heterogeneous sample.

Enrichment analysis of genome-wide association study (GWAS) gene sets pertaining to immunological disease reveals node-level differences in disease manifestation (Fig.4d). Disease-relevant gene sets are mainly found to be enriched in nodes at deeper levels of the tree, suggesting that disease manifestations are cell-type specific, rather than common features across many cells. GWAS genes relating to SLE are enriched primarily in node 22, which corresponds to topic 7. This node also is related to B cell function (Fig.4a,c), which is in line with the fact that SLE is often characterized by abnormal autoreactive B cells [19]. Node 28, corresponding to topic 13, is enriched in gene sets relating to other autoimmune diseases, such as multiple sclerosis and type-1 diabetes, as well as memory Treg cells, naive CD4+ T cells, and NK cells. Autoreactive T cell function is the primary driver of these diseases [20, 21] and the corresponding topic 13 is enriched in individuals with SLE (Fig.3d), suggesting an additional role of autoreactive T cells in SLE pathogenesis.

## 3 Discussion

### Bayesian sparsity with a structured hierarchical model

We present `LaRCH`, an embedded topic model with a built-in latent topic hierarchy. Spike-and-slab prior informed gene embedding model parameters provide a natural method of marker gene selection and interpretable latent feature detection. From multiple tests on realistic simulated scRNA-seq datasets, we show that `LaRCH` is effective at cell-type discrimination in heterogeneous samples. By enforcing an underlying topic structure, we allow for an intuitive representation of nested relationships between cell types with shared lineage.

### Insights into mechanisms of autoimmune disease

Training `LaRCH` on single-cell gene expression data across multiple disease conditions shows hierarchical genomic features in relation to disease status. Our findings show cell-type specific differences in gene expression in individuals with SLE as well as overarching cellular features across all cell-types, suggesting altered immune function. Further investigation into specific latent node-level marker genes is a promising avenue for the discovery of novel insights into the genomic drivers of autoimmunity.

### Limitations

There are a number of potential limitations to the `LaRCH` model. Using a fixed binary tree structure may lead over-parameterization of the model where many components of the tree are unused, resulting in an ill-fitting tree structure to represent the data. Unfortunately including a tree-fitting or pruning component to the model would greatly increase computational burden. We combat over-fitting in our model by implementing Bayesian priors that encourage model sparsity.

Training of the model necessitates intensive computational resources. We implemented `LaRCH` with a machine-learning library (PyTorch [22]), which often has specialized requirements for hardware, such as GPUs with sufficient processing capabilities and memory capacity. Smaller datasets of 2,000 cells require on average 12 minutes to complete model training with a tree depth of 5 in 1,000 epochs. However, on a large dataset of 1.2 million cells, training requires over 60 hours to complete.

Granularity of cell type detection proves to be a challenge with `LaRCH`. We found that rarer cell sub-types with large overlapping gene-expression profiles are difficult to discern from one another. Training a model with a deeper latent tree did not prove to yield better results.

### Future directions

The `LaRCH` model is readily expandable to fit experimental study needs. Integration of multi-omic data to include transcription factor binding affinity (ChIP-seq) and DNA accessibility profiles (ATAC-seq) is a natural addition to this model and would provide valuable insights into genomic mechanistic drivers of cell-type differentiation and disease pathology. To understand overall trends in up and down-regulated gene sets, one could relax element-wise sparsity in favour of set-wise sparsity and fine-mapping of genes, building upon the SuSiE method of variable selection techniques outline in Wang *et al*. [23]. Such novel formulation will likely result in a more scalable iterative coordinate-wise algorithm while assigning high probability mass on causal/anchor features for each independent effect. In fact, implementing a tree-structured latent feature space is a promising extension to less computationally intensive factorization-based learning algorithms [24]. Such algorithms are able to yield low-dimensional embedding of genomic data, maintaining many use cases of an embedded topic model in an efficient manner.

## 4 Methods

### 4.1 Model description

#### 4.1.1 Data generative process

We use an Embedded Topic Model (ETM) [14] approach built on a Variational Auto-Encoder (VAE) [9, 25, 26] framework borrowed from neural network architectures. To apply the topic model, each cell is treated as a document, the set of genes in the dataset as the vocabulary of size *G*, and each scRNA-seq read as a token of a gene from the vocabulary. We represent each cell as a mixture of *T* latent topics in our model.

In a classic ETM, each topic has a corresponding distribution over the gene space *β ∈* ℝ^*T ×G*^ [14]. We alter this by introducing an additional latent tree node layer to the embedding process. *K* latent nodes lie in a perfect binary tree structure of depth *D*, where *K* = 2^*D*^*−* 1. Each of the *T* latent topics, where *T* = 2^*D−*1^, corresponds to a path from the root node to a leaf node of the perfect binary tree. Each tree node now has an embedding over the gene space *β∈* ℝ^*K×V*^ and each topic also has an embedding made up of the tree nodes along its corresponding path 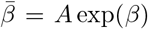, where *A* is a *T × K* matrix that captures the tree structure. From this, two topics with similar paths in the tree-structured node will share gene embeddings while the paths are identical, then differ once the paths split.

The formal data-generating process for each cell *i* in the scRNA-seq dataset is:

1. Draw latent topic proportion *θ*_*i*_ for cell *i* from:

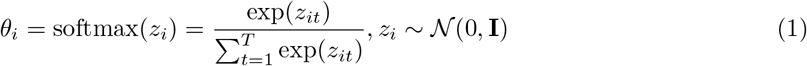
2. Determine the categorical distribution *ρ*_*i*_ of genes for cell *i* based on *θ*_*i*_:

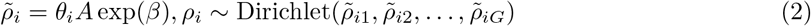
3. Draw gene *g* for each read of a cell from a multinomial distribution to give total gene counts over *R* reads:

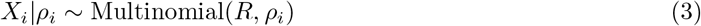

This gives the probability distribution 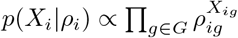

Here *z*_*i*_ is the 1 *× T* latent topic embedding of cell *i, θ*_*i*_ is the 1 *× T* topic mixture of cell *i* where 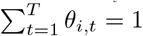, *β* is a *K* × *G* gene embedding matrix for latent nodes, *A* is a *T × K* matrix representing the tree-structure of nodes, *ρ*_*i*_ is the 1*×G* cell specific distribution of gene reads, and *X*_*i*_ is the 1*×G* count vector of genes.

#### 4.1.2 Gene embedding prior distribution

Since scRNA-seq data is typically sparse, we assume that a majority of gene embedding values *β*_*k,g*_ are statistically zero with some prior probability 1 *− π*. To promote sparsity within the model parameters, we introduce a spike-and-slab prior on the gene embedding values.

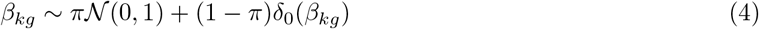

A bias parameter *b*_*g*_ is introduced to capture gene-level effects that are invariant across all topics.

#### 4.1.3 Variational inference

Since exact inference of the posterior probability of latent topics is computationally intractable in high dimensional spaces, stochastic variational inference is used as a scalable approach to finding approximate distributions [9]. We use the inference scheme as described in Kingma *et al*. [9] implemented in Pytorch using torch.autograd [22, 27]. Full variational inference and model training methods can be found in Zhang *et al*. [28].

### 4.2 Data simulation

Single-cell immune cell gene expression data was simulated using bulk-sequencing profiles from the DICE database [16]. The simulation scheme used is as follows:

1. For a given noise level *ρ* and read depth *R*
2. Determine a null and cell type-specific gene expression distribution *π*_0_ and 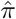, respectively

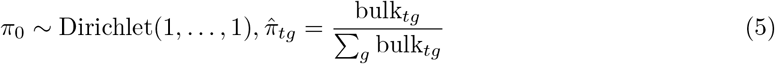

Where bulk_*tg*_ is the bulk expression profile for cell type *t* at gene *g*.
3. For a cell *i*:
  a. Randomly sample cell type *t* according to distribution *m ∼* Dirichlet(1, …, 1)
  b. Gene expression distribution for cell type *t*:

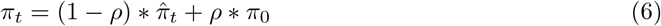
  c. Sample reads *X*_*i*_:

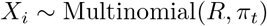

### 4.3 Marker gene detection and gene set enrichment analysis

Significant marker genes for each topic were obtained in R by feeding 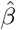 and 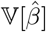 values directly into the ashr package which uses an Empirical Bayes approach for large-scale hypothesis testing and false discovery rate (FDR) estimation [29]. Significant genes are selected as those with qvalue *<* 0.05.

Ranked gene set enrichment analysis in R was performed using the fgsea R package [30] with 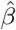 as the score and scoreType = “pos”. Gene sets used were obtained from the DICE database [16] and the NHGRI-EBI Catalog of human genome-wide association studies [31].

## 5 Data and code availability

The SLE study single-cell RNA-seq data used in this analysis can be found at GSE174188.

Code containing the most up-to-date version of the `LaRCH` package, data simulation scheme, and documentation can be found at https://github.com/causalpathlab/LaRCH

